# A molecular mechanism for LINC complex branching by structurally diverse SUN-KASH 6:6 assemblies

**DOI:** 10.1101/2020.03.21.001867

**Authors:** Manickam Gurusaran, Owen R. Davies

## Abstract

The LINC complex mechanically couples cytoskeletal and nuclear components across the nuclear envelope to fulfil a myriad of cellular functions, including nuclear shape and positioning, hearing and meiotic chromosome movements. The canonical model of the LINC complex is of individual linear nucleocytoskeletal linkages provided by 3:3 interactions between SUN and KASH proteins. Here, we provide crystallographic and biophysical evidence that SUN-KASH is a constitutive 6:6 complex in which two SUN trimers interact back-to-back. A common SUN-KASH topology is achieved through structurally diverse 6:6 interaction mechanisms by distinct KASH proteins, including zinc-coordination by Nesprin-4. The SUN-KASH 6:6 complex is incompatible with the current model of a linear LINC complex and instead suggests the formation of a branched LINC complex network. In this model, SUN-KASH 6:6 complexes act as nodes for force distribution and integration between adjacent SUN and KASH molecules, enabling the coordinated transduction of large forces across the nuclear envelope.

## Introduction

The nuclear envelope partitions nuclear components from the cytoskeleton, thereby necessitating their mechanical coupling across the nuclear envelope to enable cytoskeletal function in the structure and positioning of nuclear contents. This is achieved by the Linker of Nucleoskeleton and Cytoskeleton (LINC) complex, which traverses the nuclear envelope and binds to cytoskeletal and nuclear structures to mediate force transduction between these partitioned components (Starr & Fridolfsson, 2010) (Figure 1a). In this capacity, the LINC complex is essential for cellular life, performing critical functions in nuclear structure, shape and positioning, in addition to tissue-specific functions including sound perception in the inner ear and chromosome movements during meiosis (Lee & Burke, 2018, Meinke & Schirmer, 2015, Starr & Fridolfsson, 2010). Further, mutations of the LINC complex and its interacting partners are associated with human laminopathies, including Hutchison-Gilford progeria syndrome and Emery-Dreifuss muscular dystrophy (Meinke, Nguyen et al., 2011, Mejat & Misteli, 2010).

**Figure 1.**
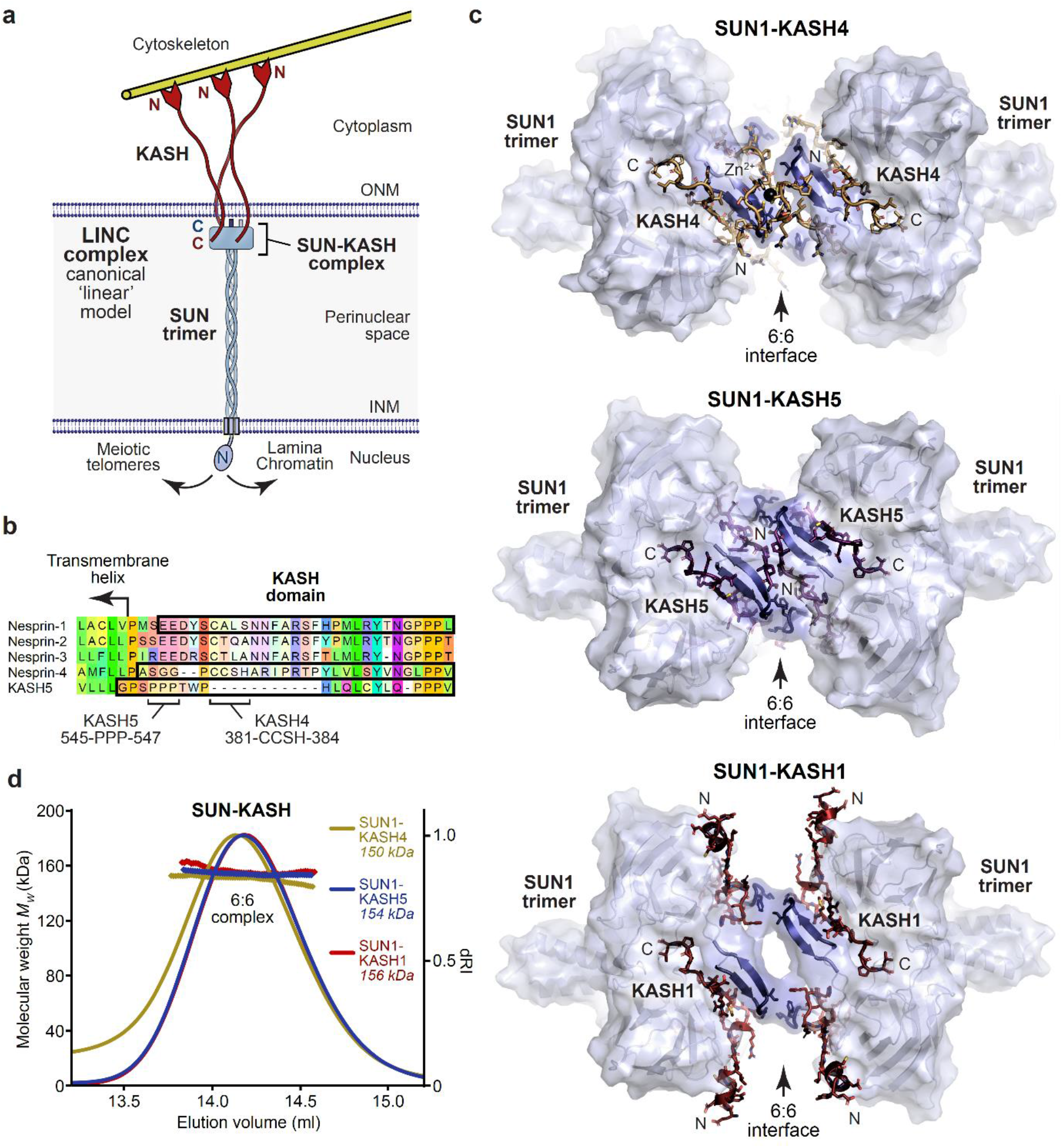
SUN-KASH complexes are 6:6 back-to-back assemblies. (**a**) The LINC complex traverses the nuclear envelope to transmit forces between the cytoskeleton and nuclear components. The canonical model of the LINC complex is a linear structure formed of SUN and Nesprin proteins, which interact via a 3:3 complex between their SUN and KASH domains within the peri-nuclear space, and cross the inner and outer nuclear membranes (INM and ONM), respectively. (**b**) Sequence alignment of the KASH domains of human Nesprins 1-4 and KASH5, highlighting the sequences used in this study (black outline), and indicating key amino-acids within KASH4 and KASH5 sequences. (**c**) Crystal structures of human SUN1-KASH4 (top), SUN1-KASH5 (middle) and SUN1-KASH1 (bottom). The SUN1 molecular surface is displayed with SUN1 KASH-lids highlighted in blue as cartoons, and KASH sequences are represented as sticks (yellow, purple and red, respectively). All structures are 6:6 complexes in which KASH proteins lie at the midline back-to-back interface between SUN1 trimers. (**d**) SEC-MALS analysis showing differential refractive index (dRI) profiles with fitted molecular weights (*Mw*) plotted as diamonds across elution peaks. SUN1-KASH4, SUN1-KASH5 and SUN1-KASH1 form 6:6 complexes in solution, with experimental molecular weights of 150 kDa, 154 kDa and 156 kDa, respectively (theoretical 6:6 – 155 kDa, 155 kDa and 157 kDa). Full elution profiles are shown in Supplementary Figure 2.

The LINC complex is formed of SUN (Sad1 and UNC84 homology) domain and KASH (Klarsicht, ANC-1 and Syne homology) domain proteins (Meinke & Schirmer, 2015, Starr & Fridolfsson, 2010), which interact immediately below the outer nuclear membrane, through complex formation between their C-terminal eponymous SUN and KASH domains (Sosa, Rothballer et al., 2012, Wang, Shi et al., 2012, Zhou, Du et al., 2012). SUN proteins then traverse the approximately 50 nm peri-nuclear space and cross the inner nuclear membrane, enabling their N-termini to bind to nuclear contents, including reported interactions with the nuclear lamina (Crisp, Liu et al., 2006, Haque, Lloyd et al., 2006, Haque, Mazzeo et al., 2010), chromatin (Chi, Haller et al., 2007) and the telomeric ends of meiotic chromosomes (Shibuya, Ishiguro et al., 2014). Similarly, KASH domain proteins cross the outer nuclear membrane and have large cytoplasmic extensions to enable their N-termini to bind to the cytoskeleton (Spindler, Redolfi et al., 2019, Starr & Fridolfsson, 2010). Thus, the LINC complex axis is established by a peri-nuclear SUN-KASH core interaction and mechanically couples the cytoskeleton and nuclear contents (Figure 1a).

In mammals, there are five SUN proteins, of which SUN1 and SUN2 are widely expressed and perform partially redundant functions (Lei, Zhang et al., 2009, Zhang, Lei et al., 2009). There are similarly multiple KASH proteins, four of which are Nesprins (Nuclear Envelope Spectrin Repeat proteins). Nesprin-1 and Nesprin-2 are widely expressed, perform overlapping functions and contain large cytoplasmic spectrin-repeat domains with N-termini that bind to actin (Banerjee, Zhang et al., 2014, Starr & Fridolfsson, 2010). Nesprin-3 shares a similar KASH domain but its cytoplasmic region binds to plectin, mediating interactions with intermediate filaments (Ketema & Sonnenberg, 2011). The two most divergent KASH proteins, Nesprin-4 and KASH5, exhibit substantial sequence diversity within their KASH domains (Figure 1b). Nesprin-4 functions in the outer hair cells of the inner ear and is essential for hearing (Horn, Brownstein et al., 2013a). Its N-terminus interacts with kinesin, which mediates microtubule binding and plus-end directed movements that achieve the basal positioning of nuclei (Horn et al., 2013a, Roux, Crisp et al., 2009). KASH5 functions in meiosis and is essential for fertility (Horn, Kim et al., 2013b, Morimoto, Shibuya et al., 2012). Its N-terminus interacts with dynein-dynactin (Horn et al., 2013b, Morimoto et al., 2012), which mediates microtubule binding and minus-end directed motility that drives rapid chromosomal movements to facilitate homologous chromosome pairing (Lee, Horn et al., 2015, Zetka, Paouneskou et al., 2020). Thus, KASH proteins execute a range of LINC complex functions in transmitting actin forces, plus-/minus-end directed microtubule movements and the tensile strength of intermediate filaments into the nucleus.

The canonical model of the LINC complex is based on crystal structures of the SUN-KASH domain complexes formed between SUN2 and Nesprin-1/2 (Sosa et al., 2012, Wang et al., 2012). The SUN domain adopts a ‘three-leaf clover’-like structure, in which a globular trimer extends from a short N-terminal trimeric coiled-coil (Sosa, Kutay et al., 2013). KASH domains are intertwined between SUN protomers and their path is defined by three distinct regions. The KASH C-terminus contains a triple proline motif that packs between the globular cores of SUN protomers. The KASH mid-region winds around the trimeric arc and is wedged between the globular core of one SUN protomer and a β-turn-β loop, known as the KASH-lid, of the adjacent protomer. The KASH N-terminus then turns by >90° to radiate out from the trimer axis and forms a disulphide bond with a SUN protomer (between SUN2 and KASH1 amino-acids C563 and C8774, respectively), which is predicted to enhance the tensile strength of SUN-KASH (Jahed, Shams et al., 2015, Sosa et al., 2012). The biological unit of SUN-KASH was interpreted from the crystal lattice as a 3:3 complex, comprising three KASH domains bound to a single SUN trimer (Sosa et al., 2012, Wang et al., 2012). On this basis, it was proposed that the LINC complex consists of a SUN-KASH 3:3 complex that is orientated vertically to allow KASH proteins to cross the outer nuclear membrane and SUN to form an extended trimeric coiled-coil that spans the peri-nuclear space (Sosa et al., 2013, Sosa et al., 2012) (Figure 1a). However, whilst analytical ultracentrifugation, SEC-MALS and gel filtration experiments have demonstrated that luminal SUN2 is trimeric (Jahed, Vu et al., 2018b, Nie, Ke et al., 2016, Sosa et al., 2012, Zhou et al., 2012), the assumed 3:3 stoichiometry of SUN-KASH complexes has not been demonstrated in solution. Further, although it has been recognised that branching or higher-order assembly between LINC complexes may be advantageous in distributing large forces and achieving coordinated motions (Jahed, Fadavi et al., 2018a, Lu, Gotzmann et al., 2008, Sosa et al., 2013, Wang et al., 2012, Zhou et al., 2012), we have hitherto lacked structural evidence in favour of this hypothesis.

Here, we provide crystallographic and biophysical evidence in support of the LINC complex forming a branched network. We find that SUN-KASH complexes between SUN1 and Nesprin-4, KASH5 and Nesprin-1 are 6:6 structures in which KASH-binding induces constitutive back-to-back interactions between SUN trimers. The three distinct KASH domains provide structurally diverse but related 6:6 interfaces that achieve the same topology with potential hinge-like motion between SUN trimers. The SUN-KASH 6:6 complex is incompatible with the current model of a linear LINC complex, and instead suggests the formation of a branched LINC complex network. Thus, we propose that SUN-KASH domain complexes act as nodes for branching and integration between LINC complexes to achieve the coordinated transduction of large forces across the nuclear envelope.

## Results

### SUN-KASH complexes are 6:6 hetero-oligomers

The previously reported crystal structures of SUN-KASH complexes between SUN2 and Nesprins 1-2 revealed almost identical structures that were interpreted as 3:3 hetero-oligomers (Sosa et al., 2012, Wang et al., 2012). The KASH domains of Nesprin-4 and KASH5 exhibit sequence divergence from Nesprins 1-3, including the presence of N-terminal motifs of 381-CCSH-384 and 545-PPP-547, which are conserved within Nesprin-4 and KASH5 sequences, respectively (Figure 1b). On this basis, we reasoned that Nesprin-4 and KASH5 may impose unique SUN-KASH structures that differ from the classical architecture of Nesprin 1-3 complexes, which may underlie their specialised functional roles. We thus solved the X-ray crystal structures of SUN-KASH complexes formed between the SUN domain of SUN1 and KASH domains of Nesprin-4 and KASH5 (herein referred to as SUN1-KASH4 and SUN1-KASH5). The SUN1-KASH4 structure was solved at a resolution of 2.75 Å and revealed a 6:6 assembly in which two globular 3:3 complexes are held in a back-to-back configuration through zinc-coordination by opposing KASH4 molecules across the 6:6 interface (Figure 1c, Table 1 and Supplementary Figure 1a,b). The SUN1-KASH5 crystal structure was solved at 1.54 Å resolution and revealed a similar 6:6 assembly in which opposing 3:3 complexes are held together by extensive interactions between opposing KASH5 molecules and KASH-lids (Figure 1c, Table 1 and Supplementary Figure 1a,c). Thus, both Nesprin-4 and KASH5 form SUN-KASH 6:6 hetero-oligomers in which similar topologies of back-to-back 3:3 complexes are achieved through structurally diverse 6:6 interfaces.

**Table 1.**
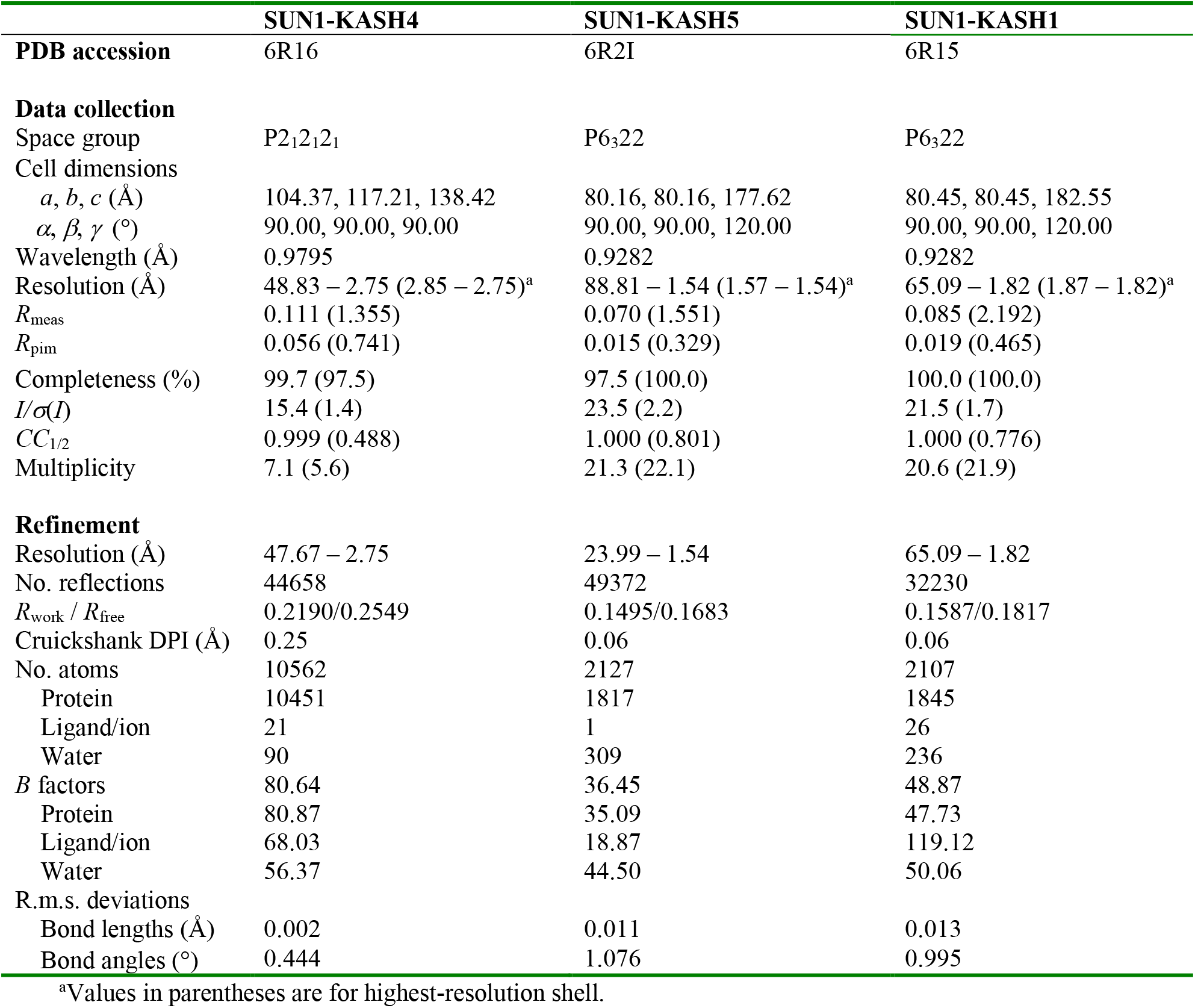
Data collection, phasing and refinement statistics.

Is the 6:6 assembly unique to SUN-KASH complexes formed by Nesprin-4 and KASH5? We next solved the crystal structure of the SUN-KASH complex between SUN1 and Nesprin-1 (herein referred to as SUN1-KASH1). The SUN1-KASH1 structure was solved at 1.82 Å resolution and demonstrated a similar 6:6 back-to-back assembly, albeit with less extensive interface-spanning interactions provided solely by opposing KASH-lids (Figure 1c, Table 1 and Supplementary Figure 1a,d). The electron density indicated the presence of a molecule bound close to the 6:6 interface, which we interpreted as a disordered HEPES molecule from the crystallisation condition. This likely provided structural rigidity that underlies the high resolution of the dataset, but was not essential for the structure as we solved numerous other datasets at lower resolution in which an identical 6:6 interface was present in absence of a bound molecule (data not shown). Importantly, the SUN1-KASH1 structure closely matches the previous SUN2-KASH1/2 structures, in which similar 6:6 interfaces were present in the crystal lattice but were dismissed as crystal contacts (Supplementary Figure 1e) (Sosa et al., 2012, Wang et al., 2012). It was thus critical to determine whether SUN1-KASH1 is a 6:6 complex in solution. We utilised size-exclusion chromatography multi-angle light scattering (SEC-MALS) as the gold standard for determining molecular species. SEC-MALS revealed that all SUN-KASH complexes exist solely as 6:6 hetero-oligomers (Figures 1d and 2a, and Supplementary Figure 2). Moreover, their 6:6 complexes remained intact at the lowest detectable concentrations (Figure 2b-d) and we failed to detect 3:3 complexes in any biochemical conditions tested. Thus, we conclude that SUN-KASH complexes are 6:6 hetero-oligomers in which two 3:3 complexes interact in back-to-back configurations. The SUN-KASH 6:6 structure is incompatible with the current model of a linear 3:3 LINC complex and instead demonstrates a constitutive interaction between SUN trimers that could mediate the physical coupling of adjacent LINC complexes within the peri-nuclear space.

**Figure 2.**
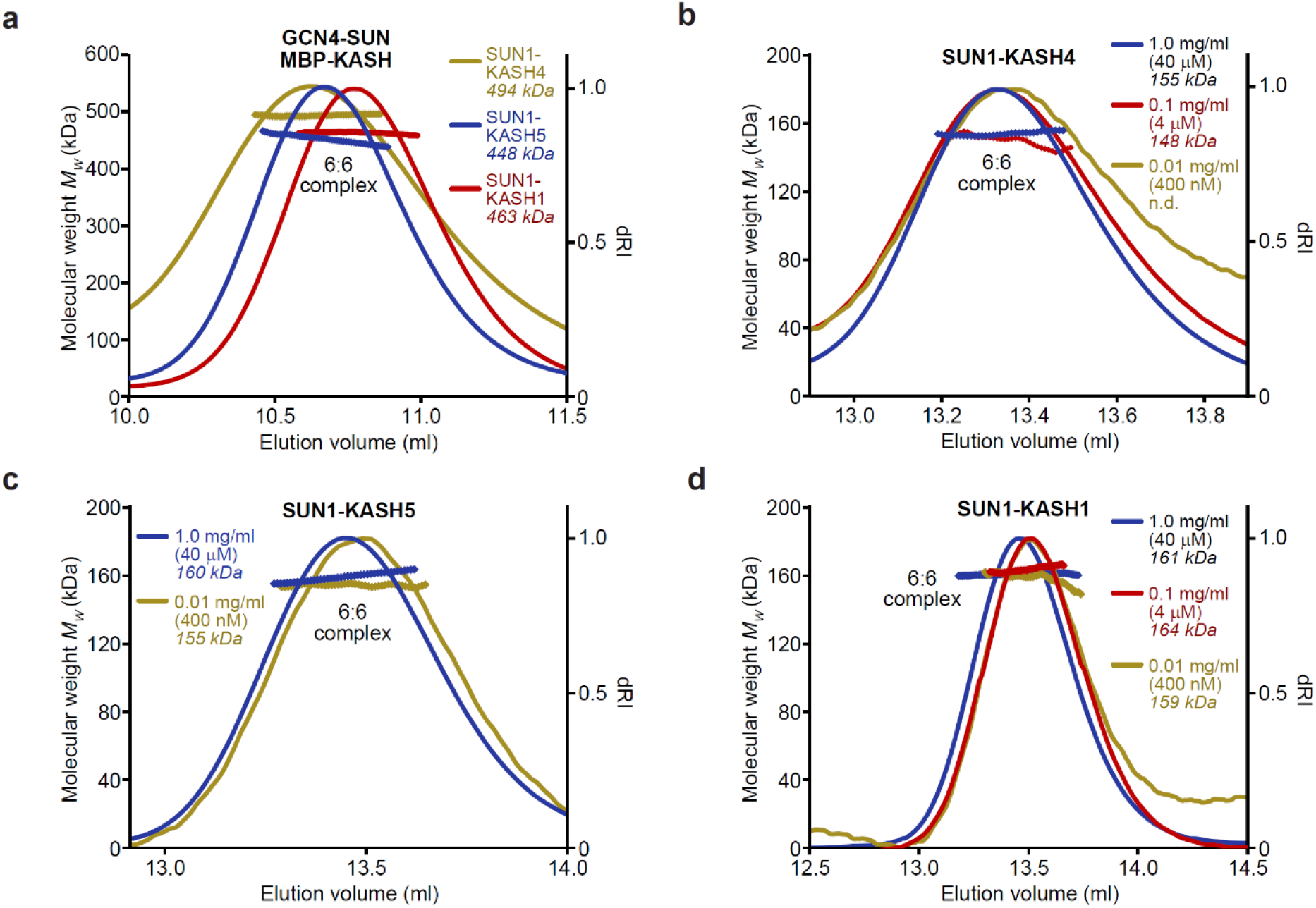
SUN-KASH 6:6 complexes are stable in solution. (**a-d**) SEC-MALS analysis. (**a**) GCN4-SUN1 and MBP-KASH form 6:6 complexes of 494 kDa (KASH4, yellow), 448 kDa (KASH5, blue) and 463 kDa (KASH1, red) (theoretical 6:6 – 464 kDa, 464 kDa and 466 kDa). (**b-d**) Dilution series of SUN-KASH complexes analysed at 1.0 mg/ml (blue), 0.1 mg/ml (red) and 0.01 mg/ml (yellow) for (**b**) SUN1-KASH4 (theoretical 6:6 - 155 kDa), (**c**) SUN1-KASH5 (theoretical 6:6 - 155 kDa) and (**d**) SUN1-KASH1 (theoretical 6:6 - 157 kDa).

### Structural diversity within the SUN-KASH 6:6 interface

Our SUN-KASH crystal structures reveal the formation of similar 6:6 architectures through diverse back-to-back interfaces. Whilst the C-termini of all three KASH domains adopt the same structure, their N-termini differ substantially (Figure 3a,b). KASH1 undergoes a turn of >90° to radiate from the trimer axis, similar to the previously reported SUN2-KASH1/2 structures (Supplementary Figure 1e), whereas KASH4 and KASH5 follow the arc of the SUN1 trimer, enabling them to contribute directly to the 6:6 interface (Figure 3a,b).

**Figure 3.**
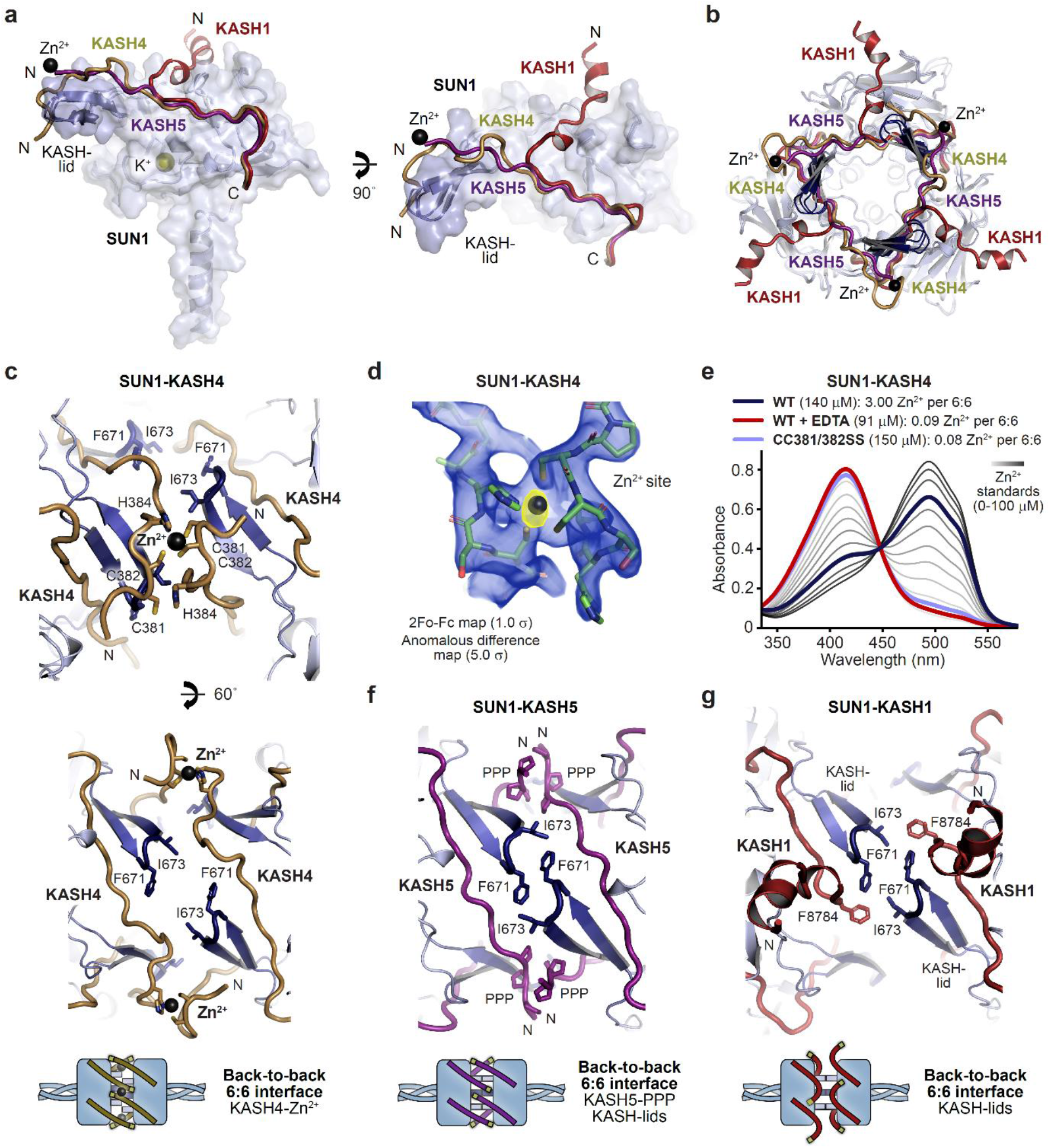
Specialised KASH sequences provide distinct SUN-KASH 6:6 assembly mechanisms. (**a**) SUN-KASH 1:1 protomers from SUN1-KASH4, SUN1-KASH5 and SUN1-KASH1 crystal structures, superposed, and displayed as the SUN1 molecular surface with KASH-lids highlighted in blue as cartoons, and KASH sequences represented as cartoons (yellow, purple and red, respectively). (**b**) Cross-section through the back-to-back interface of superposed SUN1-KASH4, SUN1-KASH5 and SUN1-KASH1 6:6 assemblies such that their constituent 3:3 complexes are visible. (**c**) Structural details of the SUN1-KASH4 6:6 interface, showing a zinc-binding site in which opposing KASH4 chains provide asymmetric ligands C381 and C382, and C382 and H384 (top), and the lack of interface-spanning interactions between opposing SUN1 KASH-lids (bottom). (**d**) 2Fo-Fc (blue) and anomalous difference (yellow) electron density maps contoured at 1.0 σ and 5.0 σ, respectively, at a zinc-binding site of SUN1-KASH4. (**e**) Spectrophotometric determination of zinc content for SUN1-KASH4 wild-type (dark blue; 3.00 Zn^2+^ per 6:6), wild-type with EDTA treatment prior to gel filtration (red; 0.09 Zn^2+^ per 6:6), and CC381/382SS (light blue; 0.08 Zn^2+^ per 6:6), using metallochromic indicator PAR, with zinc standards shown in a gradient from light to dark grey (0-100 μM). (**f**) Structural details of the SUN1-KASH5 6:6 interface, demonstrating interface-spanning interactions between PPP-motifs (amino-acids 545-PPP-547) of opposing KASH5 chains, and between amino-acids F671 and I673 of opposing SUN1 KASH-lids. (**g**) Structural details of the SUN1-KASH1 6:6 interface showing interactions between amino-acids F671 and I673 of opposing SUN1 KASH-lids that are supported by KASH1 amino-acid F8784, but with no interface-spanning interactions between opposing KASH1 chains.

SUN1-KASH4 adopts an unusual conformation in which the 6:6 complex is held together by three interface spanning zinc-sites, each coordinated by opposing KASH4 molecules (Figure 3c and Supplementary Figure 3a). The presence of metal ions in the crystal structure was confirmed by corresponding peaks in anomalous difference electron density maps (Figure 3d), and their identity as zinc ions that were co-purified from bacterial expression was confirmed by the spectrophotometric determination of three zinc ions per 6:6 complex in solution that were lost upon pre-incubation with EDTA (Figure 3e). The zinc-sites are coordinated by asymmetric ligands from 381-CCSH-384 motifs of opposing KASH4 molecules, comprising C381 and C382 from one molecule, and C382 and H384 from the other (Figure 3c), and mutation of both cysteine residues to serine was sufficient to preclude zinc-binding (Figure 3e). The three zinc-sites form a tripod of interactions that provide the sole interface-spanning contacts between opposing 3:3 complexes (Figure 3b).

SUN1-KASH5 demonstrates the most extensive 6:6 interface in which KASH5 molecules and SUN1 KASH-lids from opposing 3:3 complexes wind around each other in a right-handed screw to create a complete circumferential interface enclosing a hollow core, similar to a β-barrel fold (Figure 3f and Supplementary Figure 3b). KASH5 follows an almost linear path, packed between a SUN1 globular core and KASH-lids of opposing SUN1 protomers, with N-terminal 545-PPP-547 motifs of opposing molecules interacting across the interface. KASH5 and KASH4 follow similar paths, with KASH5 PPP-motif interactions and KASH4 zinc-sites located at the same positions and providing analogous interface-spanning interactions (Figure 3a,b). However, an important distinction is that a torsional rotation of approximately 20° between the 3:3 complexes of SUN1-KASH5, relative to SUN1-KASH4, brings together opposing KASH-lids and enables their interaction across the interface (Figure 3f). Thus, tip-to-tip interactions via amino-acids I673 and F671 of opposing SUN1 KASH-lids contribute to the extensive 6:6 interface of SUN1-KASH5 (Figure 3f).

The SUN1-KASH1 6:6 complex is formed solely of a tripod of KASH-lid tip-to-tip interactions mediated by amino-acids I673 and F671, in the same manner and owing to the same torsional rotation as in the SUN1-KASH5 structure (Figure 3g and Supplementary Figure 3c). KASH1 undergoes acute angulation away from the 6:6 interface (Figure 3a,b), as previously observed in SUN2-KASH1/2 (Supplementary Figure 1e). As such, whilst amino-acid F8784 binds to the KASH-lids of each tip-to-tip interaction site (Figure 3g), KASH1’s N-terminus does not contribute to the 6:6 interface (Figure 3a,b). This creates an open interface, with large solvent channels between opposing 3:3 complexes (Supplementary Figure 3c). Overall, the three structures demonstrate alternative SUN-KASH 6:6 interaction mechanisms that are differentially exploited by KASH proteins. SUN1-KASH4 and SUN1-KASH1 represent extreme states of this spectrum in which the 6:6 interface is mediated solely by KASH-mediated metal coordination and SUN1 KASH-lids, respectively. In contrast, SUN1-KASH5 adopts an intermediate structure that utilises both KASH and KASH-lid interaction mechanisms to form a fully enclosed 6:6 interface.

### SUN1-KASH1 complex formation depends on KASH-lid 6:6 interactions

On the basis of our SUN-KASH crystal structures, we predicted that KASH-lid tip-to-tip interactions are essential for 6:6 hetero-oligomer formation in solution by SUN1-KASH1 but not SUN1-KASH4. We tested this by generating glutamate mutations of KASH-lid tip amino-acids I673 and F671, which mediate interface-spanning tip-to-tip interactions within SUN1-KASH1 and SUN1-KASH5 but have no contacts within their respective 3:3 complexes (Figure 3c,f,g and Supplementary Figure 4a-c). We also analysed a glutamate mutation of amino-acid W676, which mediates hydrophobic interactions with the KASH domain within a constituent 3:3 complex (Supplementary Figure 4a-c), and acted as a negative control in disrupting all three SUN-KASH complexes (Figure 4a,b). It was not possible to analyse SUN-KASH binding through amylose pull-down owing to the non-specific binding between SUN1 and amylose resin (Supplementary Figure 4e). Instead, we exploited this phenomenon by using amylose resin to purify complexes and dissociated proteins following GCN4-SUN1 and MBP-KASH co-expression, which we enriched by ion exchange (Supplementary Figure 4d), and then pooled all fractions containing SUN-KASH complexes and dissociated proteins for analysis by analytical gel filtration (Figure 4a,b). We validated the resulting elution profiles through SEC-MALS by confirming that the wild-type fusion complexes and dissociated GCN4-SUN1 and MBP-KASH proteins are 6:6 complexes, trimers and monomers, respectively (Supplementary Figure 4f).

**Figure 4.**
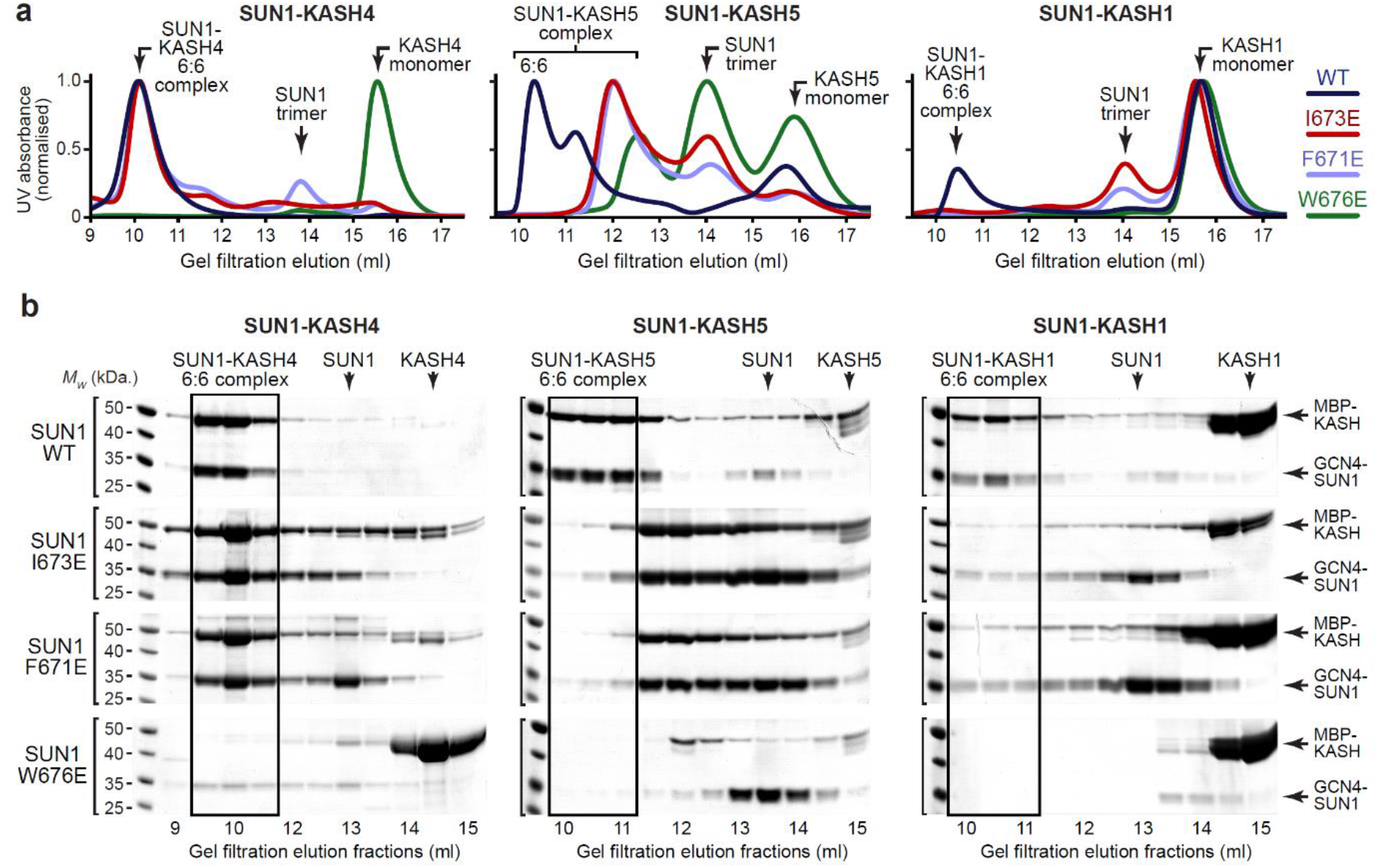
SUN1 KASH-lid residues involved in 6:6 assembly are essential for KASH1-binding. (**a,b**) Gel filtration analysis. GCN4-SUN1 and MBP-KASH proteins were co-expressed and purified by amylose affinity (utilising non-specific binding by SUN1 for non-interacting mutants) and ion exchange (Supplementary Figure 4d), and all fractions containing SUN-KASH complexes and dissociated proteins were concentrated and loaded onto an analytical gel filtration column. The elution profiles were validating by SEC-MALS in which wild-type fusion complexes and dissociated GCN4-SUN1 and MBP-KASH1 proteins were found to be 6:6 complexes, trimers and monomers, respectively (Supplementary Figure 4f). (**a**) Gel filtration chromatograms (UV absorbance at 280 nm) across elution profiles for SUN1 wild-type (WT; dark blue), I673E (red), F671E (light blue) and W676E (green), with KASH4 (left), KASH5 (middle) and KASH1 (right), and (**b**) SDS-PAGE of their corresponding elution fractions. Source data are provided as a Source Data file

The SUN1-KASH4 6:6 complex was impervious to KASH-lid mutations I673E and F671E (Figure 4a,b and Supplementary Figures 4d and 5a), in keeping with the lack of KASH-lid tip-to-tip interactions at its 6:6 interface. In stark contrast, SUN1-KASH1 was disrupted by I673E and F671E mutations (Figure 4a,b and Supplementary Figure 4d), confirming that KASH-lid tip-to-tip interactions are essential for its 6:6 complex formation. Upon removal of the trimerizing GCN4 tag, the dissociated SUN1 I673E protein was monomeric, similar to wild-type SUN1 expressed in absence of a KASH domain (Supplementary Figure 5b), and SAXS analysis confirmed that its SUN domain remained folded (Supplementary Figure 5c-g and Table 2). The failure to observe smaller hetero-oligomers demonstrates that SUN1-KASH1 3:3 complexes are unstable in absence of the 6:6 interface, indicating that SUN1-KASH1 is a constitutive 6:6 hetero-oligomer.

**Table 2.** Summary of SEC-SAXS data.

|  | SUN1<br>I673E<br>(monomer) | SUN1-KASH4<br>(6:6) | SUN1-KASH1<br>(6:6) | SUN1-KASH5<br>(6:6) |
| --- | --- | --- | --- | --- |
| **Guinier analysis** |  |  |  |  |
| $I(0)$ (cm<sup>-1</sup>) | 0.042 | 0.045 | 0.130 | 0.010 |
| $R_g$ (Å) | 21 | 40 | 39 | 38 |
| $q_{min}$ (Å<sup>-1</sup>) | 0.0080 | 0.0014 | 0.0090 | 0.0070 |
| **$P(r)$ analysis** |  |  |  |  |
| $I(0)$ (cm<sup>-1</sup>) | 0.042 | 0.045 | 0.132 | 0.10 |
| $R_g$ (Å) | 22 | 40 | 39 | 39 |
| $D_{max}$ (Å) | 82 | 135 | 130 | 135 |
| Porod volume (Å<sup>3</sup>) | 39367 | 292301 | 303602 | 274824 |
| MW from Porod volume (kDa) | 23 | 172 | 179 | 162 |
| $V_c$ (Å<sup>2</sup>) | 238 | 825 | 853 | 784 |
| MW from $V_c$ (kDa) | 22 | 139 | 152 | 131 |
| **DAMMIF *ab initio* modelling**<br>**(30 models)** |  |  |  |  |
| Symmetry | P1 | N/A | N/A | N/A |
| NSD mean | 0.645 | N/A | N/A | N/A |
| $\chi^2$ (reference model) | 1.85 | N/A | N/A | N/A |
| **Structural modelling** |  |  |  |  |
| CRY SOL - crystal structure ($\chi^2$) | 5.43 | 1.62 | 4.83 | 5.50 |
| CORAL - modelling of N-termini ($\chi^2$) | N/A | 1.25 | 4.55 | 1.70 |
| CORAL - rigid body modelling ($\chi^2$) | N/A | N/A | 1.56 | N/A |
| SREFLEX - normal mode analysis ($\chi^2$) | 1.72 – 1.98 | N/A | N/A | N/A |

In agreement with the equal roles of KASH domain and KASH-lid interactions at its 6:6 interface, SUN1-KASH5 exhibited intermediate phenotypes upon I673E and F671E mutation, with retention of complex formation but reduction in oligomer size to species that likely reflect partially dissociating 6:6 complexes (Figure 4a,b and Supplementary Figure 4d). We conclude that the diverse roles of KASH-lids at the 6:6 interfaces of SUN-KASH crystal structures are truly reflective of their solutions states and that KASH-lid tip-to-tip interactions are essential for assembly of a constitutive SUN1-KASH1 6:6 hetero-oligomer.

### Hinge-like motion of the SUN-KASH 6:6 interface

How could the SUN-KASH 6:6 complex be orientated within the nuclear envelope? Its back-to-back assembly suggests a horizontal orientation, parallel to the outer nuclear membrane, with SUN trimers organised obliquely within the peri-nuclear space. In this configuration, tension forces carried by SUN and KASH molecules would exert bending moments on the structure, favouring a hinge-like angulation between opposing 3:3 complexes. We thus utilised small-angle X-ray scattering (SAXS) to determine whether SUN-KASH complexes adopt angled conformations in solution. Whilst SAXS data of SUN1-KASH4 and SUN1-KASH5 were closely fitted by their crystal structures upon flexible modelling of missing termini (χ^2^ values of 1.25 and 1.70), we achieved only poor fits for SUN1-KASH1 (χ^2^ = 4.83) (Figure 5a,b and Supplementary Figure 6 and Table 2). In case of large-scale motion, we performed SAXS-based rigid-body modelling using two SUN1-KASH1 3:3 complexes as independent rigid bodies. We consistently obtained models that closely fitted experimental data (χ^2^ = 1.56) in which 3:3 complexes interact back-to-back with a bend of approximately 60° relative to the crystal structure (Figure 5c,d and Table 2). In this model, two pairs of KASH-lid tip-to-tip interactions by I673 and F671 are retained, whilst the third is disrupted, and an additional interface is formed between opposing central KASH-lids. Thus, KASH-lids act as a hinge at the 6:6 interface, opening the linear crystal structure into an angled conformation that is seemingly more stable in solution.

**Figure 5.**
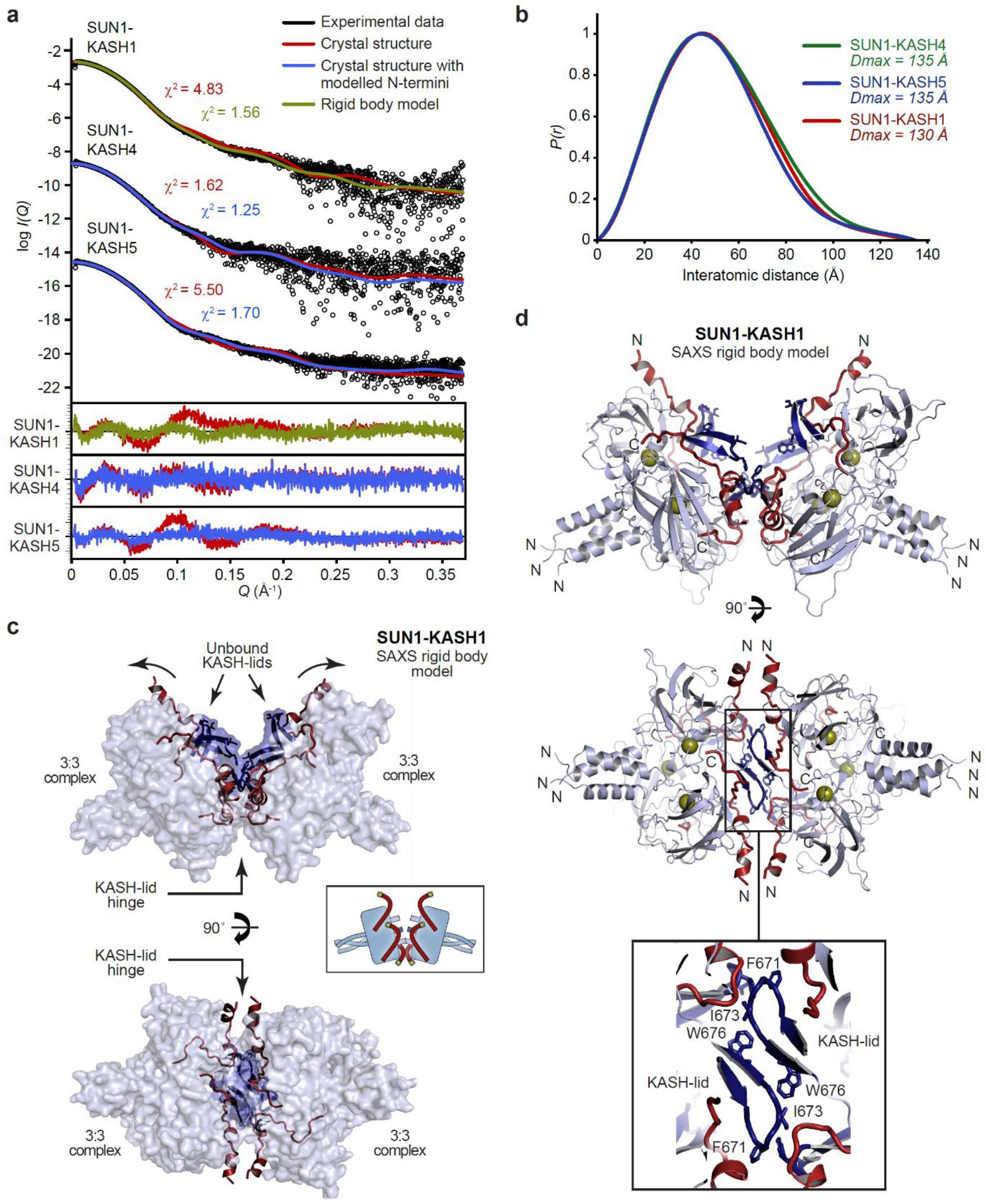
SEC-SAXS analysis of SUN-KASH 6:6 complexes. (**a**) SAXS scattering curves of SUN1-KASH4, SUN1-KASH5 and SUN1-KASH1 overlaid with theoretical scattering curves of their crystal structures (red), crystal structures with KASH flexible N-termini modelled by CORAL (blue) and rigid body model of two 3:3 complexes (yellow). Residuals for each fit are shown (inset). (**b**) SAXS *P*(*r*) distributions showing maximum dimensions of 135 Å, 135 Å and 130 Å, respectively. (**c-d**) SAXS rigid body model of SUN1-KASH1 shown as (**c**) surface and (**d**) cartoon representation, in which two constituent 3:3 complexes from its crystal structure were assigned as rigid bodies, with the 6:6 assembly generated by fitting to experimental SAXS data of solution SUN1-KASH1 (χ^2^ = 1.56). (**d**) The cartoon representation highlights structural details of the predicted KASH-lid interface, including the presence of unbound KASH-lids, and the close approximation of opposing KASH-lids, which achieve an asymmetric positioning of the N-termini of KASH domains in locations and orientations compatible with their upstream sequences crossing the outer nuclear membrane.

The hinged SUN1-KASH1 structure solves a critical problem in understanding the potential role of the 6:6 complex within its cellular context. Whilst the linear crystal structure distributes the KASH1 N-termini around its circumferential exterior (Figures 1c and 3b), making it difficult to envisage how all KASH1 molecules could access the outer nuclear membrane, the asymmetrical hinged structure places all six KASH1 N-termini in favourable positions and orientations for their upstream transmembrane sequences to cross the outer nuclear membrane (Figure 5c,d).

Is a similar hinge-like angulation possible for SUN1-KASH4 and SUN1-KASH5? Whilst their extensive 6:6 interfaces retain linear structures in solution (Figure 5a-b, Supplementary Figure 6 and Table 2), angulation may be achieved by tension forces. We thus performed normal mode analysis to determine whether angled structures are conformationally accessible. We observed low frequency normal modes corresponding to hinge-like angulation at the 6:6 interface for all SUN-KASH complexes (Figure 6), indicating that angled conformations are accessible flexible states. As described for SUN1-KASH1, hinging of SUN1-KASH4 and SUN-KASH5 would place the N-termini of their constituent KASH domains in suitable positions and orientations to cross the outer nuclear membrane, so adoption of hinged conformations may be a critical part of forming stable membrane-associated assemblies. We thus conclude a model in which hinged SUN-KASH 6:6 complexes, parallel with the outer nuclear membrane, act as nodes for the integration and distribution of tension forces between oblique SUN trimers and KASH molecules within a branched LINC complex network (Figure 7).

**Figure 6.**
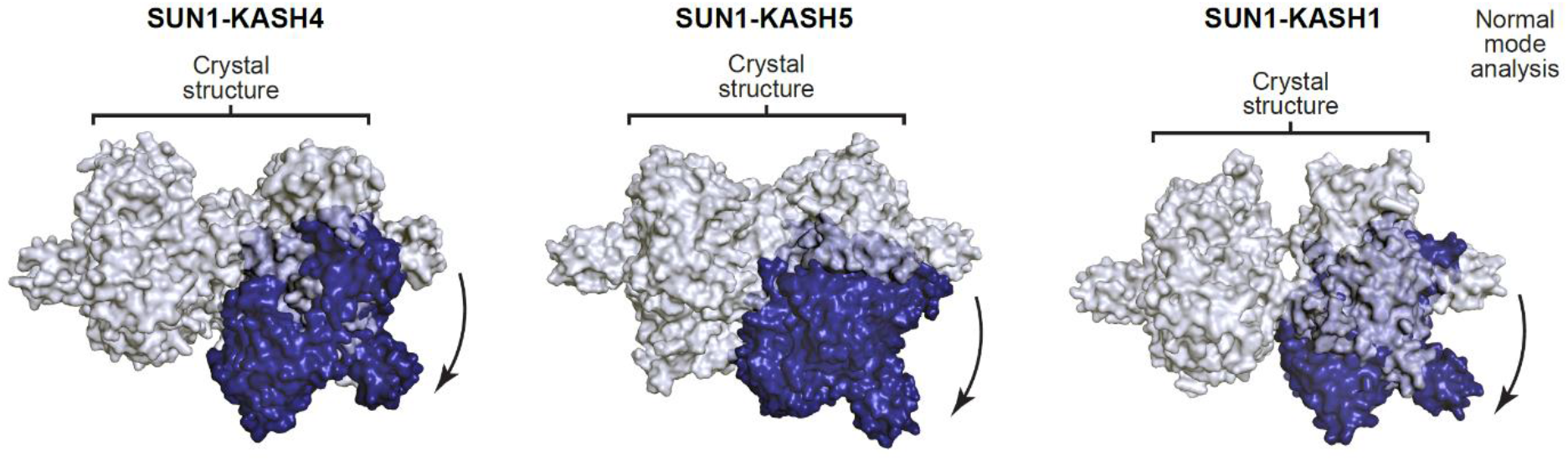
Hinge-link conformational flexibility within SUN-KASH 6:6 assemblies. Normal mode analysis of SUN-KASH complexes in which non-linear normal modes calculated by the NOLB algorithm are shown as the largest amplitude of motion of one constituent 3:3 complex (blue) relative to its original position and its stationary opposing 3:3 complex within the crystal structure (grey) for SUN1-KASH4 (top), SUN1-KASH5 (middle) and SUN1-KASH1 (bottom).

**Figure 7.**
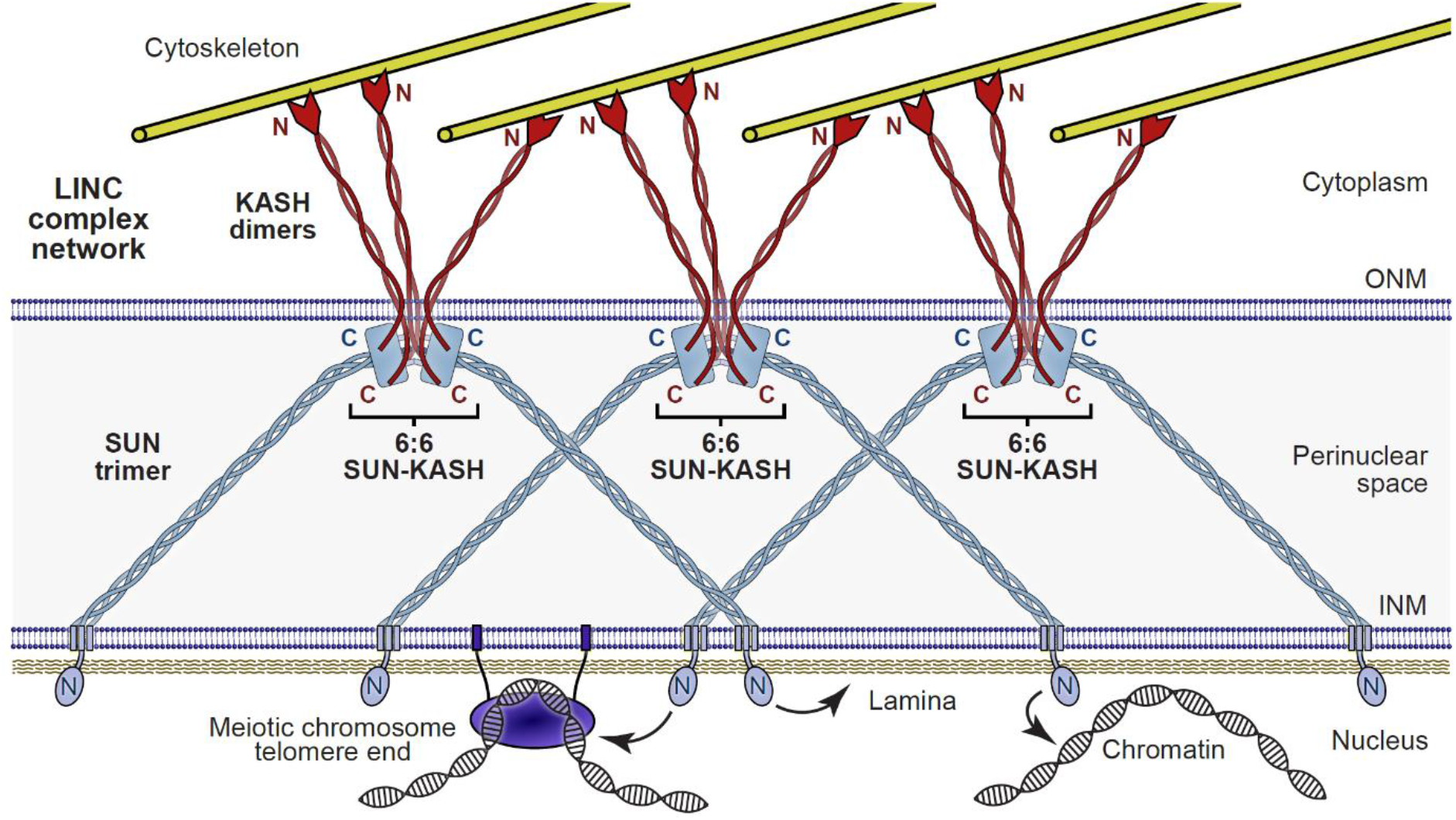
The LINC complex as a branched network of SUN-KASH assemblies. Model of the LINC complex as a branched network in which SUN-KASH 6:6 complexes act as nodes for force integration and distribution between two SUN trimers and three KASH dimers, which can bind to spatially-separated and distinct nuclear and cytoskeletal components, respectively. This model enables cooperation between adjacent molecules within a LINC complex network to facilitate the transduction of large and coordinated forces across the nuclear envelope.

## Discussion

How does our finding of a constitutive SUN-KASH 6:6 assembly integrate with previous studies of the LINC complex? It has been shown by analytical ultracentrifugation, SEC-MALS and gel filtration studies that luminal SUN2 is trimeric, with its isolated SUN domain being trimeric or monomeric, depending on biochemical conditions (Jahed et al., 2018b, Sosa et al., 2013, Wang et al., 2012, Zhou et al., 2012). These findings agree with our observations that the isolated SUN domain of SUN1 is a monomer that undergoes trimerization upon addition of an N-terminal coiled-coil. The only previous analysis of SUN-KASH *in vitro* was through demonstrating complex formation by analytical gel filtration, without means for oligomer determination (Esra Demircioglu, Cruz et al., 2016). Thus, the canonical 3:3 model was generated by combining the known trimerization of SUN proteins with a single trimer of the SUN2-KASH1/2 crystal structure, without evidence of whether this represents its true biological unit in solution. The SEC-MALS and SEC-SAXS analyses reported herein provide the first reported evidence of the SUN-KASH solution structure. In this, we determined that SUN-KASH complexes exist solely as the 6:6 hetero-oligomers observed in our SUN1-KASH1/4/5 crystal structures and that are also present in the crystal lattices of previous SUN2-KASH1/2 structures (Sosa et al., 2012, Wang et al., 2012). Importantly, we did not detect 3:3 complexes in any conditions tested. Hence, our conclusion that SUN-KASH complexes are constitutive 6:6 structures is consistent with, and provides the only feasible explanation for, all existing crystallographic, biochemical and biophysical data.

How does the SUN-KASH 6:6 assembly relate to LINC complex structure and function within the cell? Whilst it is not currently possible to determine the oligomeric state of LINC complexes *in situ* within the cell, we propose that SUN-KASH 6:6 assembly mediates LINC complex branching within the nuclear envelope. A branched network is attractive as it can transmit forces of magnitudes far in excess of those carried by single molecules, is impervious to disruption of individual linkages, and allows communication and coordination between adjacent molecules. Thus, SUN-KASH 6:6 hetero-oligomers could enable force distribution and integration between adjacent molecules to achieve coordinated actions within the nucleus. Importantly, LINC complex branching by SUN-KASH 6:6 assembly is compatible with all existing cellular data, and moreover explains a number of observations of higher-order LINC complex structures, including its immobility within the nuclear envelope (Lu et al., 2008), foci formation within the meiotic nuclear envelope (Ding, Xu et al., 2007, Horn et al., 2013b, Morimoto et al., 2012), and the formation of transmembrane actin-associated nuclear (TAN) lines (Luxton, Gomes et al., 2010). Nevertheless, given the myriad of LINC complex functions in almost all eukaryotic cells (Lee & Burke, 2018, Meinke & Schirmer, 2015, Starr & Fridolfsson, 2010), and the dynamics of nuclear envelope formation and dissolution during the cell cycle, we recognise the possibility of multiple LINC complex conformations. Indeed, SUN1 and SUN2 adopt autoinhibitory conformations in absence of KASH-binding (Nie et al., 2016, Xu, Li et al., 2018), which likely represent unassembled states, and SUN-KASH complexes could conceivably interact with other nuclear envelope components to stabilise alternative conformations. Thus, whilst our proposed branched LINC complex is consistent with all existing data, we acknowledge that alternative LINC complex conformations may exist within the spatial and temporal contexts of disparate cell types.

The back-to-back nature of SUN-KASH 6:6 complexes suggests their orientation parallel to the outer nuclear membrane, with SUN trimers organised obliquely within the peri-nuclear space (Figure 7). Our SAXS analysis of SUN1-KASH1 indicates that it adopts a hinged conformation in solution, stabilised by two KASH-lid tip-to-tip interactions and the lateral association of central KASH-lids. This provides a more extensive interface than the tripod of KASH-lid tip-to-tip interactions within the crystal structure, explaining its preferential formation outside of a crystal lattice. In contrast, SUN1-KASH4 and SUN1-KASH5 have extensive 6:6 interfaces that preserve linearity in solution. Nevertheless, normal mode analysis predicted that hinged structures of all three SUN-KASH complexes are conformationally accessible states. As such, SUN-KASH 6:6 hetero-oligomers may become angled in response to tension forces carried by obliquely orientated SUN and KASH molecules, which would position the N-termini of all six KASH domains in an appropriate orientation and proximity for their upstream transmembrane sequences to cross the outer nuclear membrane. Thus, the hinged conformation could be mutually reinforced by tension forces between oblique SUN and KASH molecules, and the steric restrictions imposed by the proximity between SUN-binding KASH domains and their upstream transmembrane sequences. Further, all three SUN-KASH 6:6 interfaces consist of circumferential exteriors and hollow cores that are largely hydrophobic in nature. Thus, we envisage that the SUN-KASH hinged structure may be stabilised by interaction of its 6:6 interface with phospholipids of the outer nuclear membrane, potentially forming an integrated membrane-bound complex between a 6:6 hetero-oligomer and the transmembrane regions of its six KASH proteins.

How do the diverse 6:6 interfaces provided by KASH proteins facilitate their specialised functions? An intriguing observation is that Nesprin-4 and KASH5, which transduce microtubule forces (Horn et al., 2013a, Horn et al., 2013b, Morimoto et al., 2012, Roux et al., 2009), demonstrate extensive interactions at their 6:6 interfaces. In contrast, a far less extensive 6:6 interface is found in classical Nesprins, which transduce actin forces and the tensile strength of intermediate filaments (Banerjee et al., 2014, Ketema & Sonnenberg, 2011, Starr & Fridolfsson, 2010). Thus, cytoskeletal components may have differential requirements for the strength, structure and stability of SUN-KASH 6:6 hetero-oligomers. Further, their diverse interfaces may provide disparate mechanisms for the timely assembly and disassembly of LINC complex networks during miotic and meiotic cell cycles. This could be achieved by inhibition of the 6:6 interface, which is sufficient to disrupt the SUN1-KASH1 globular complex. Further, as the SUN1-KASH4 6:6 interface is mediated by zinc-coordination sites, its formation could be regulated by zinc availability and sequestration within the peri-nuclear space.

The SUN-KASH 6:6 hetero-oligomer provides the first structural evidence, but perhaps not the sole structural means, of LINC complex network formation. Additional branching events could result from variation in oligomer state along the SUN-KASH axis. Whilst SUN is assumed to form a trimeric coiled-coil extending from its globular core, its protein sequence is inconsistent with formation of a single continuous coiled-coil. Instead, it suggests the presence of short discrete coiled-coils, of potentially distinct oligomers, that are connected by unstructured linkers. Further, C526 residues of SUN1 are reported to form disulphide bonds (Lu et al., 2008), suggesting tethering between SUN trimers from separate SUN-KASH 6:6 complexes. Thus, oligomer variation and disulphide formation may combine with the SUN-KASH 6:6 interface to provide multiple branching events within the peri-nuclear space. Whilst there is no reported structural information regarding KASH proteins, we find that KASH5 is dimeric (Manickam & Davies, unpublished findings), indicating branching between SUN-KASH 6:6 complexes and their cytoskeletal attachments. Thus, we propose that coordinated force transduction is achieved by a highly branched LINC complex network. In this model, the lowest degree of branching supported by current structural evidence is of SUN-KASH 6:6 hetero-oligomers mediating force distribution and integration between three KASH dimers and two SUN trimers (Figure 7).

## Methods

### Recombinant protein expression and purification

The SUN domain of human SUN1 (amino-acid residues 616-812) was fused to an N-terminal TEV-cleavable His_6_-GCN4 tag (as described in Sosa et al. (2012)) and cloned into a pRSF-Duet1 (Merck Millipore) vector. The KASH domains of human KASH5 (amino-acid residues 542-562), Nesprin-4 (KASH4, amino-acid residues 376-404) and Nesprin-1 (KASH1, amino-acid residues 8769-8797) were cloned into pMAT11 (Peranen, Rikkonen et al., 1996) vectors for expression as TEV-cleavable His_6_-MBP fusion proteins respectively. SUN1 and KASH constructs were co-expressed in BL21 (DE3) cells (Novagen®), in 2xYT media, induced with 0.5 mM IPTG for 16 hours at 25°C. Cell disruption was achieved by sonication in 20 mM Tris pH 8.0, 500 mM KCl and cellular debris removed by centrifugation at 40,000 g. Fusion proteins were purified through consecutive Ni-NTA (Qiagen), amylose (NEB), and HiTrap Q HP (GE Healthcare) ion exchange chromatography. TEV protease was utilised to remove affinity tags and cleaved samples were purified through ion exchange chromatography and size exclusion chromatography (HiLoad^™^ 16/600 Superdex 200, GE Healthcare) in 20 mM Tris pH 8.0, 150 mM KCl, 2 mM DTT. Protein samples were concentrated using Microsep^™^ Advance Centrifugal Devices 10,000 MWCO centrifugal filter units (PALL) and were stored at −80°C following flash-freezing in liquid nitrogen. Protein samples were analysed by SDS-PAGE and visualised with Coomassie staining. Concentrations were determined by UV spectroscopy using a Cary 60 UV spectrophotometer (Agilent) with extinction coefficients and molecular weights calculated by ProtParam (http://web.expasy.org/protparam/).

### Crystal structure solution of SUN1-KASH4 (PDB accession 6R16)

SUN1-KASH4 protein crystals were obtained through vapour diffusion in sitting drops, by mixing 100 nl of protein at 25 mg/ml with 100 nl of crystallisation solution (0.06 M MgCl_2_; 0.06 M CaCl_2_; 0.1 M Imidazole pH 6.5; 0.1M MES (acid) pH 6.5; 18% Ethylene glycol; 18% PEG 8K) and equilibrating at 20°C for 4-9 days. Crystals were flash frozen in liquid nitrogen. X-ray diffraction data were collected at 0.9795 Å, 100 K, as 2000 consecutive 0.10° frames of 0.040 s exposure on a Pilatus 6M-F detector at beamline I04 of the Diamond Light Source synchrotron facility (Oxfordshire, UK). Data were indexed and integrated in XDS (Kabsch, 2010), scaled in XSCALE (Diederichs, McSweeney et al., 2003) and merged using Aimless (Evans, 2011). Crystals belong to orthorhombic spacegroup P212121 (cell dimensions a = 104.37 Å, b = 117.21 Å, c = 138.42 Å, α = 90°, β = 90°, γ = 90°), with six copies of SUN1 and KASH4 per asymmetric unit. The structure was solved by molecular replacement using Phaser (McCoy, Grosse-Kunstleve et al., 2007), with SUN1-KASH1 (this study, PDB accession 6R15) as a search model. The structure was re-built by PHENIX Autobuild (Adams, Afonine et al., 2010) and completed through iterative manual model building in Coot (Emsley, Lohkamp et al., 2010), with the addition of six potassium ions, three zinc ions and ethylene glycol ligands. The structure was refined using PHENIX refine (Adams et al., 2010) with isotropic atomic displacement parameters and TLS parameters, using SUN1-KASH1 as a reference structure. The structure was refined against 2.75 Å data to *R* and *Rfree* values of 0.2190 and 0.2549 respectively, with 98.22% of residues within the favoured regions of the Ramachandran plot (0 outliers), clashscore of 4.89 and overall MolProbity score of 1.26 (Chen, Arendall et al., 2010). The final SUN1-KASH4 model was analysed using the *Online_DPI* webserver (http://cluster.physics.iisc.ernet.in/dpi) to determine a Cruikshank diffraction precision index (DPI) of 0.25 Å (Kumar, Gurusaran et al., 2015).

### Crystal structure solution of SUN1-KASH5 (PDB accession 6R2I)

SUN1-KASH5 protein crystals were obtained through vapour diffusion in sitting drops, by mixing 100 nl of protein at 25 mg/ml with 100 nl of crystallisation solution (0.12 M 1,6-Hexanediol; 0.12 M 1-Butanol 1,2-Propanediol (racemic); 0.12 M 2-Propanol; 0.12 M 1,4-Butanediol; 0.12 M 1,3-Propanediol; 0.1 M Imidazole pH 6.5; 0.1 M MES (acid) pH 6.5; 18% Glycerol; 18% PEG 4K) and equilibrating at 20°C for 4-9 days. Crystals were flash frozen in liquid nitrogen. X-ray diffraction data were collected at 0.9282 Å, 100 K, as 2000 consecutive 0.10° frames of 0.050 s exposure on a Pilatus 6M-F detector at beamline I04-1 of the Diamond Light Source synchrotron facility (Oxfordshire, UK).

Data were indexed, integrated, scaled and merged in AutoPROC using XDS (Kabsch, 2010) and Aimless (Evans, 2011). Crystals belong to hexagonal spacegroup P6322 (cell dimensions a = 80.16 Å, b = 80.16 Å, c = 177.62 Å, α = 90°, β = 90°, γ = 120°), with one copy of SUN1 and KASH5 per asymmetric unit. The structure was solved by molecular replacement using Phaser (McCoy et al., 2007), with SUN1-KASH1 (this study, PDB accession 6R15) as a search model. The structure was re-built by PHENIX Autobuild (Adams et al., 2010) and completed through iterative manual model building in Coot (Emsley et al., 2010), with the addition of a potassium ion. The structure was refined using PHENIX refine (Adams et al., 2010), using anisotropic atomic displacement parameters. The structure was refined against 1.54 Å data to *R* and *Rfree* values of 0.1495 and 0.1683, respectively, with 96.71% of residues within the favoured regions of the Ramachandran plot (0 outliers), clashscore of 6.11 and overall MolProbity score of 1.54 (Chen et al., 2010). The final SUN1-KASH5 model was analysed using the *Online_DPI* webserver (http://cluster.physics.iisc.ernet.in/dpi) to determine a Cruikshank diffraction precision index (DPI) of 0.06 Å (Kumar et al., 2015).

### Crystal structure solution of SUN1-KASH1 (PDB accession 6R15)

SUN1-KASH1 protein crystals were obtained through vapour diffusion in sitting drops, by mixing 100 nl of protein at 21 mg/ml with 100 nl of crystallisation solution (0.09 M NaF; 0.09 M NaBr; 0.09 M NaI; 0.1M Sodium HEPES pH 7.5; 0.1 M MOPS (acid) pH 7.5; 18% PEGMME 550; 18% PEG 20K) and equilibrating at 20°C for 4-9 days. Crystals were flash frozen in liquid nitrogen. X-ray diffraction data were collected at 0.9282 Å, 100 K, as 2000 consecutive 0.10° frames of 0.100 s exposure on a Pilatus 6M-F detector at beamline I04-1 of the Diamond Light Source synchrotron facility (Oxfordshire, UK). Data were indexed, integrated, scaled and merged in Xia2 (Winter, 2010) using XDS (Kabsch, 2010), XSCALE (Diederichs et al., 2003) and Aimless (Evans, 2011). Crystals belong to hexagonal spacegroup P6_3_22 (cell dimensions a = 80.45 Å, b = 80.45 Å, c = 182.55 Å, α = 90°, β = 90°, γ = 120°), with one copy of SUN1 and KASH1 per asymmetric unit. The structure was solved by molecular replacement using Phaser (McCoy et al., 2007), with SUN2-KASH1 (PDB accession 4DXR) (Sosa et al., 2012) as a search model. The structure was re-built by PHENIX Autobuild (Adams et al., 2010) and completed through iterative manual model building in Coot (Emsley et al., 2010), with the addition of a potassium ion, and PEG and HEPES ligands. The structure was refined using PHENIX refine (Adams et al., 2010), using isotropic atomic displacement parameters with four TLS groups per chain. The structure was refined against 1.82 Å data to *R* and *Rfree* values of 0.1587 and 0.1817, respectively, with 96.86% of residues within the favoured regions of the Ramachandran plot (0 outliers), clashscore of 0.00 and overall MolProbity score of 0.69 (Chen et al., 2010). The final SUN1-KASH1 model was analysed using the *Online_DPI* webserver (http://cluster.physics.iisc.ernet.in/dpi) to determine a Cruikshank diffraction precision index (DPI) of 0.06 Å (Kumar et al., 2015).

### Size-exclusion chromatography multi-angle light scattering (SEC-MALS)

The absolute molar masses of protein samples and complexes were determined by size-exclusion chromatography multi-angle light scattering (SEC-MALS). Protein samples at >1 mg/ml (unless otherwise states) were loaded onto a Superdex^™^ 200 Increase 10/300 GL size exclusion chromatography column (GE Healthcare) in 20 mM Tris pH 8.0, 150 mM KCl, 2 mM DTT, at 0.5 ml/min using an ÄKTA^™^ Pure (GE Healthcare). The column outlet was fed into a DAWN^®^ HELEOS^™^ II MALS detector (Wyatt Technology), followed by an Optilab^®^ T-rEX^™^ differential refractometer (Wyatt Technology). Light scattering and differential refractive index data were collected and analysed using ASTRA^®^ 6 software (Wyatt Technology). Molecular weights and estimated errors were calculated across eluted peaks by extrapolation from Zimm plots using a dn/dc value of 0.1850 ml/g. SEC-MALS data are presented as differential refractive index (dRI) profiles with fitted molecular weights (*M_w_*) plotted across elution peaks.

### Spectrophotometric determination of zinc content

The presence of zinc in protein samples was determined through a spectrophotometric method using the metallochromic indicator 4-(2-pyridylazo) resorcinol (PAR) (Sabel, Shepherd et al., 2009). Protein samples at 90-200 μM, corresponding to SUN1-KASH4 wild-type and CC381/382SS, and a wild-type sample that had been treated with EDTA prior to gel-filtration, were digested with 0.6 μg/μl proteinase K (NEB) at 60°C for 1 hour. 10 μl of the supernatant of each protein digestion was added to 80 μl of 50 μM 4-(2-pyridylazo)-resorcinol (PAR) in 20 mM Tris, pH 8.0, 150 mM KCl, incubated for 5 min at room temperature, and UV absorbance spectra were recorded between 600 and 300 nm (Varian Cary 60 spectrophotometer). Zinc concentrations were estimated from the ratio between absorbance at 492 and 414 nm, plotted on a line of best fit obtained from analysis of 0–100 μM zinc acetate standards.

### KASH-binding by SUN1 point mutants

The wild-type and individual point mutations I673E, F671E and W676E of SUN1 (as His_6_-GCN4 fusions) were co-expressed with KASH (as His_6_-MBP fusion) as described above. Initial purification was performed by amylose affinity chromatography (NEB), relying on the residual affinity of SUN1 in cases when point mutations were disruptive. Resultant protein mixtures were analysed by ion exchange chromatography using HiTrap Q HP (GE Healthcare) and comparable samples from full elution profiles of wild-type and mutant proteins for each KASH binding-partner were analysed by SDS-PAGE. The entire elutions were then pooled, concentrated and analysed by size-exclusion chromatography on a Superdex^™^ 200 Increase 10/300 GL size exclusion chromatography column (GE Healthcare) in 20 mM Tris pH 8.0, 150 mM KCl, 2 mM DTT, at 0.5 ml/min using an ÄKTA^™^ Pure (GE Healthcare). Elution fractions of wild-type and mutant proteins for each KASH binding-partner were analysed by SDS-PAGE.

### Size-exclusion chromatography small-angle X-ray scattering (SEC-SAXS)

SEC-SAXS experiments were performed at beamline B21 of the Diamond Light Source synchrotron facility (Oxfordshire, UK). Protein samples at concentrations >10 mg/ml were loaded onto a Superdex^™^ 200 Increase 10/300 GL size exclusion chromatography column (GE Healthcare) in 20 mM Tris pH 8.0, 150 mM KCl at 0.5 ml/min using an Agilent 1200 HPLC system. The column outlet was fed into the experimental cell, and SAXS data were recorded at 12.4 keV, detector distance 4.014 m, in 3.0 s frames. ScÅtter 3.0 (http://www.bioisis.net) was used to subtract, average the frames and carry out the Guinier analysis for the radius of gyration (*Rg*), and *P*(*r*) distributions were fitted using *PRIMUS* (P.V.Konarev, 2003). *Ab initio* modelling was performed using *DAMMIF* (Franke & Svergun, 2009); 30 independent runs were performed in P1 and averaged. Crystal structures and models were fitted to experimental data using *CRYSOL* (Svergun D.I., 1995). Normal mode analysis was used to model conformational flexibility for fitting to SAXS data using *SREFLEX* (Panjkovich & Svergun, 2016), and rigid body and flexible termini modelling was performed using *CORAL* (Petoukhov, Franke et al., 2012).

### Normal mode analysis of SUN1-KASH structures

Non-linear normal modes were calculated and visualised for SUN1-KASH 6:6 structures using the NOLB algorithm (Hoffmann & Grudinin, 2017) within the normal mode analysis SAMSON element (https://www.samson-connect.net).

### Protein sequence and structure analysis

Nesprin sequences were aligned and visualised using MUSCLE (Madeira, Park et al., 2019) and Jalview (Waterhouse, Procter et al., 2009). Molecular structure images were generated using the PyMOL Molecular Graphics System, Version 2.3 Schrödinger, LLC.

## Data availability

Crystallographic structure factors and atomic coordinates have been deposited in the Protein Data Bank (PDB) under accession numbers 6R15, 6R16 and 6R2I, and raw diffraction data have been uploaded to https://proteindiffraction.org/. All other data are available from the corresponding author upon reasonable request. Uncropped gel images relating to Figure 4b are available in the source data file.

## Acknowledgements

We thank Diamond Light Source and the staff of beamlines I04, I04-1 and B21 (proposals mx13587, mx18598 and sm15836), A. Basle for assistance with X-ray crystallographic data collection, and J. Dunce for critically reviewing the manuscript. O.R.D. is a Sir Henry Dale Fellow jointly funded by the Wellcome Trust and Royal Society (Grant Number 104158/Z/14/Z).

## Author contributions

M.G. performed all experimental work. O.R.D. designed experiments, analysed data and wrote the manuscript.

---

**Supplementary Figure 1.**
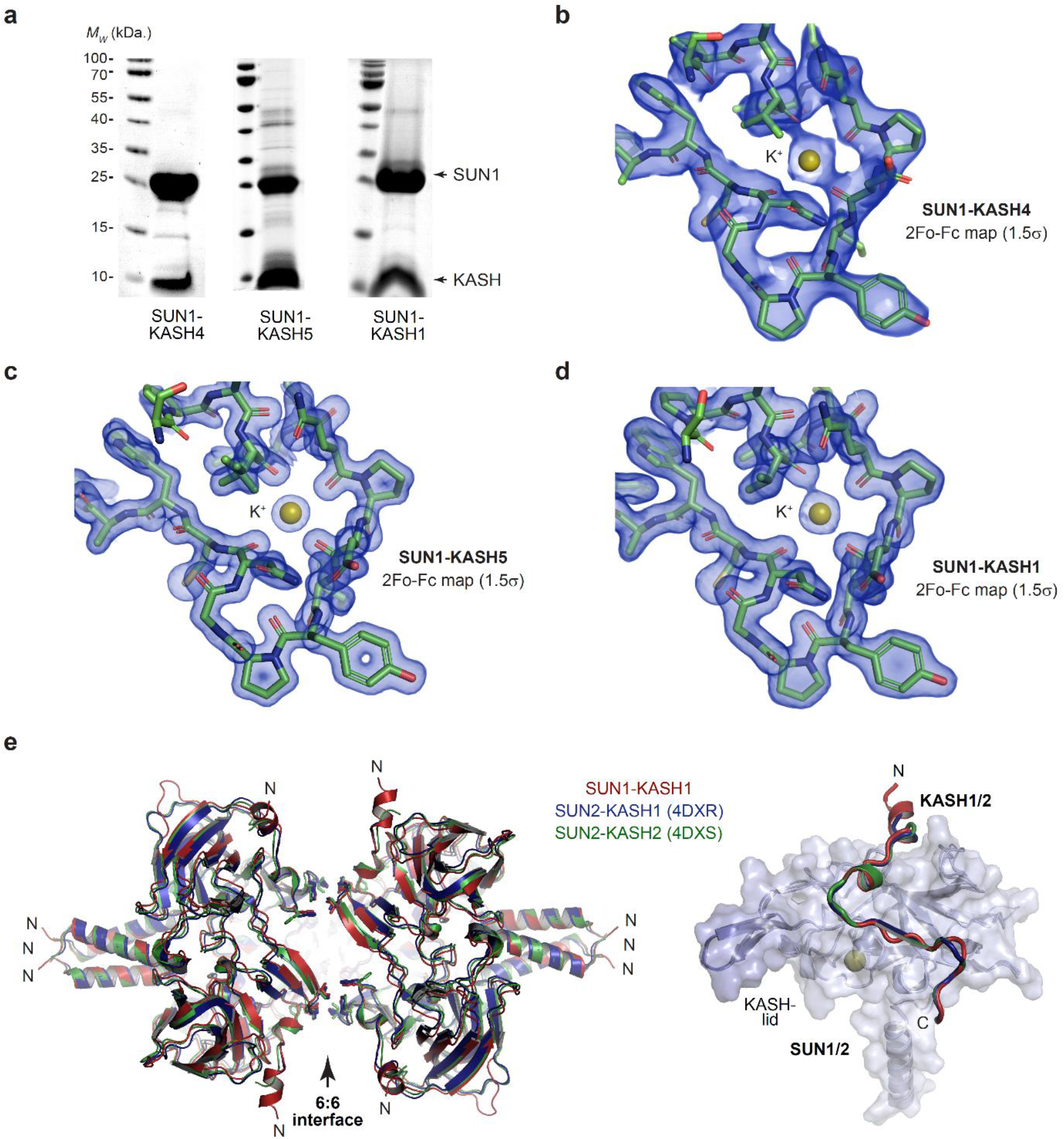
Crystal structures of SUN-KASH complexes. (**a**) SDS-PAGE of purified SUN1-KASH4 (left), SUN1-KASH5 (middle) and SUN1-KASH1 (right) complexes used for crystallographic and biophysical experiments. (**b-d**) 2Fo-Fc electron density maps contoured at 1.5 σ of the same orientation of the SUN1 K^+^-binding site for (**b**) SUN1-KASH4, (**c**) SUN1-KASH5 and (**d**) SUN1-KASH1. (**e**) Superposition of SUN1-KASH1 (red) with SUN2-KASH1 (blue; PDB accession 4DXR) and SUN2-KASH2 (green; PDB accession 4DXS) (Sosa et al., 2012), showing the common 6:6 assembly that is present within their crystal lattices (left) and their constituent 1:1 protomers (right).

**Supplementary Figure 2.**
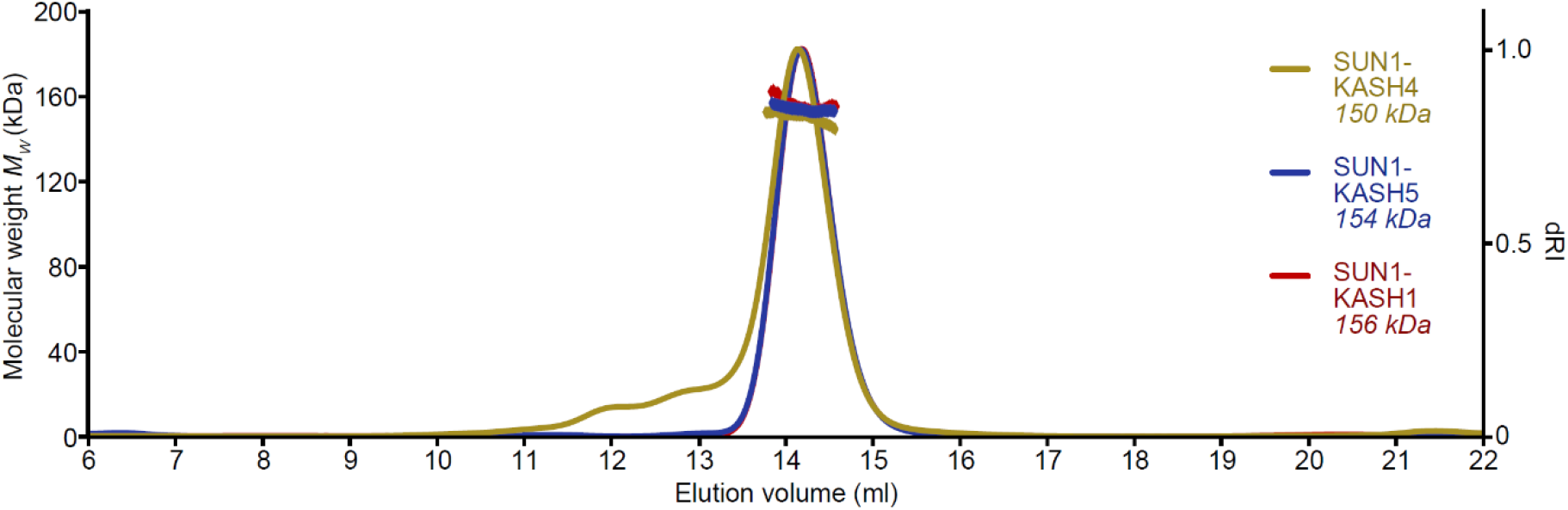
SUN-KASH complexes are 6:6 assemblies. Full SEC-MALS elution profiles corresponding to the peaks shown in Figure 1d. SUN1-KASH4, SUN1-KASH5 and SUN1-KASH1 form 6:6 complexes in solution, with experimental molecular weights of 150 kDa, 154 kDa and 156 kDa, respectively (theoretical 6:6 – 155 kDa, 155 kDa and 157 kDa).

**Supplementary Figure 3.**
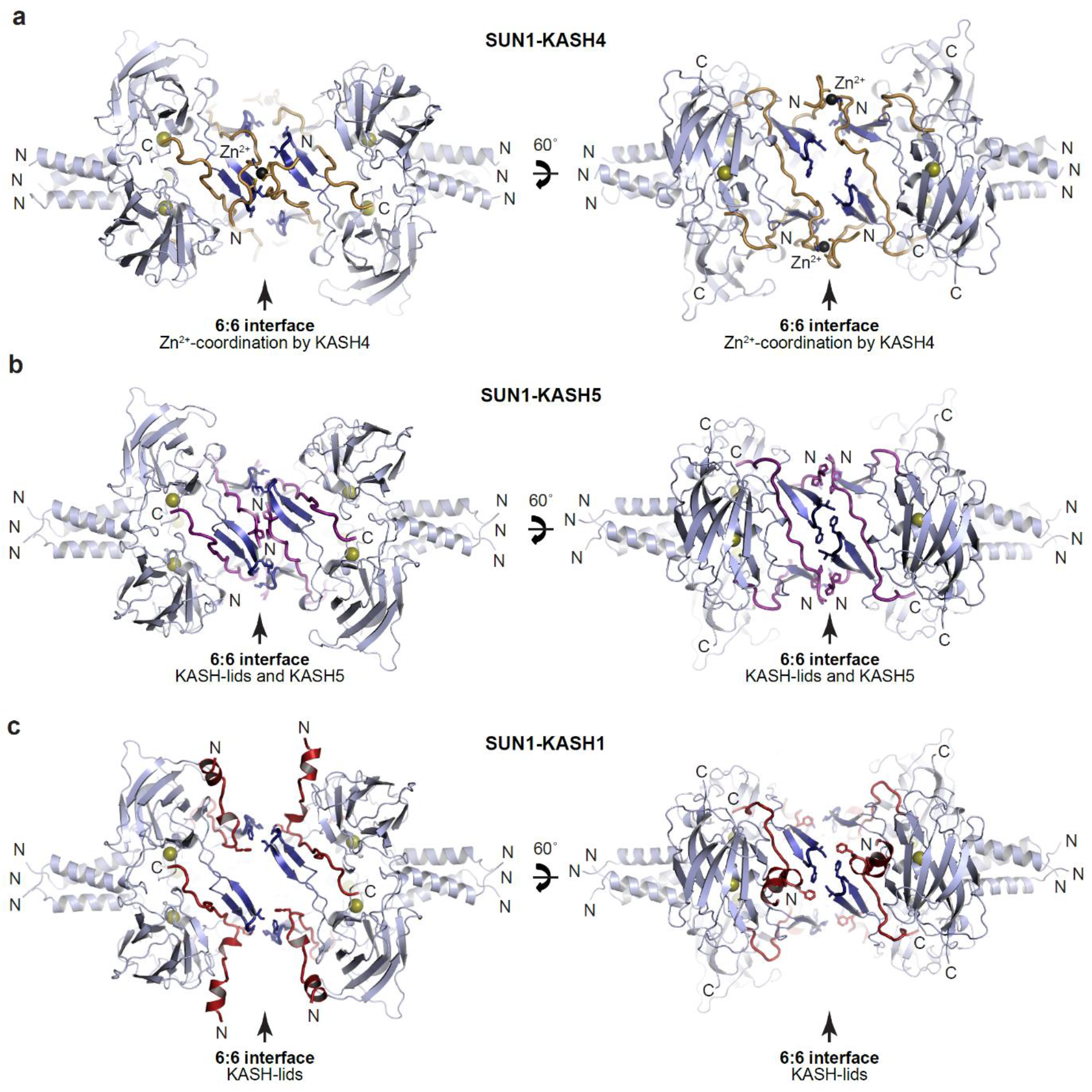
SUN-KASH crystal structures. (**a-c**) Crystal structures of (**a**) SUN1-KASH4, (**b**) SUN1-KASH5 and (**c**) SUN1-KASH1 shown in cartoon representation with SUN1 KASH-lids highlighted in blue and KASH4, KASH5 and KASH1 shown in yellow, purple and red, respectively.

**Supplementary Figure 4.**
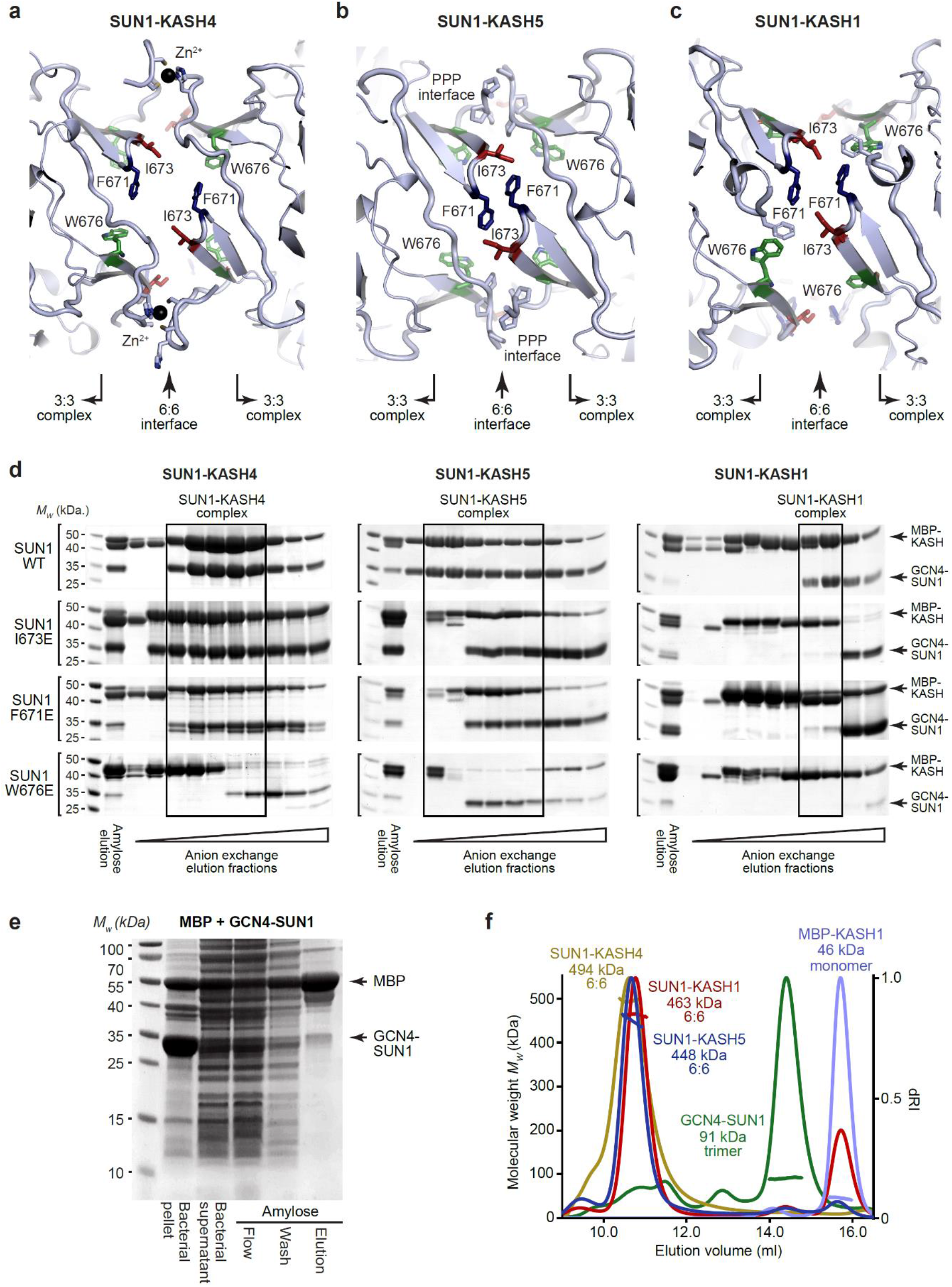
SUN-KASH complex formation upon SUN1 KASH-lid mutagenesis. (**a-c**) Structural details of the 6:6 interface of (**a**) SUN1-KASH4, (**b**) SUN1-KASH5 and (**c**) SUN1-KASH1, highlighting SUN1 KASH-lid residues I673 (red), F671 (blue) and W676 (green). (**d**) SDS-PAGE of anion exchange elution profiles of GCN4-SUN1 and MBP-KASH proteins that were co-expressed and purified by amylose affinity (utilising non-specific binding by SUN1 for non-interacting mutants), comprising SUN1 wild-type, I673E, F671E and W676E, with KASH4 (left), KASH5 (middle) and KASH1 (right). All fractions containing SUN-KASH complexes or dissociated proteins were pooled, concentrated and loaded onto an analytical gel filtration column for the analyses shown in Figure 4a,b. (**e**) Amylose pulldown following co-expression of MBP and GCN4-SUN1, showing that GCN4-SUN1 binds non-specifically to amylose resin. (**f**) SEC-MALS analysis demonstrating that SUN1-KASH1/4/5 fusion complexes, GCN4-SUN1 and MBP-KASH1 are 6:6 complexes, trimers and monomers, respectively (theoretical masses – 466 kDa, 464 kDa, 464 kDa, 88 kDa and 48 kDa). This analysis provides validation for the gel filtration elution profiles shown in Figure 4a,b.

**Supplementary Figure 5.**
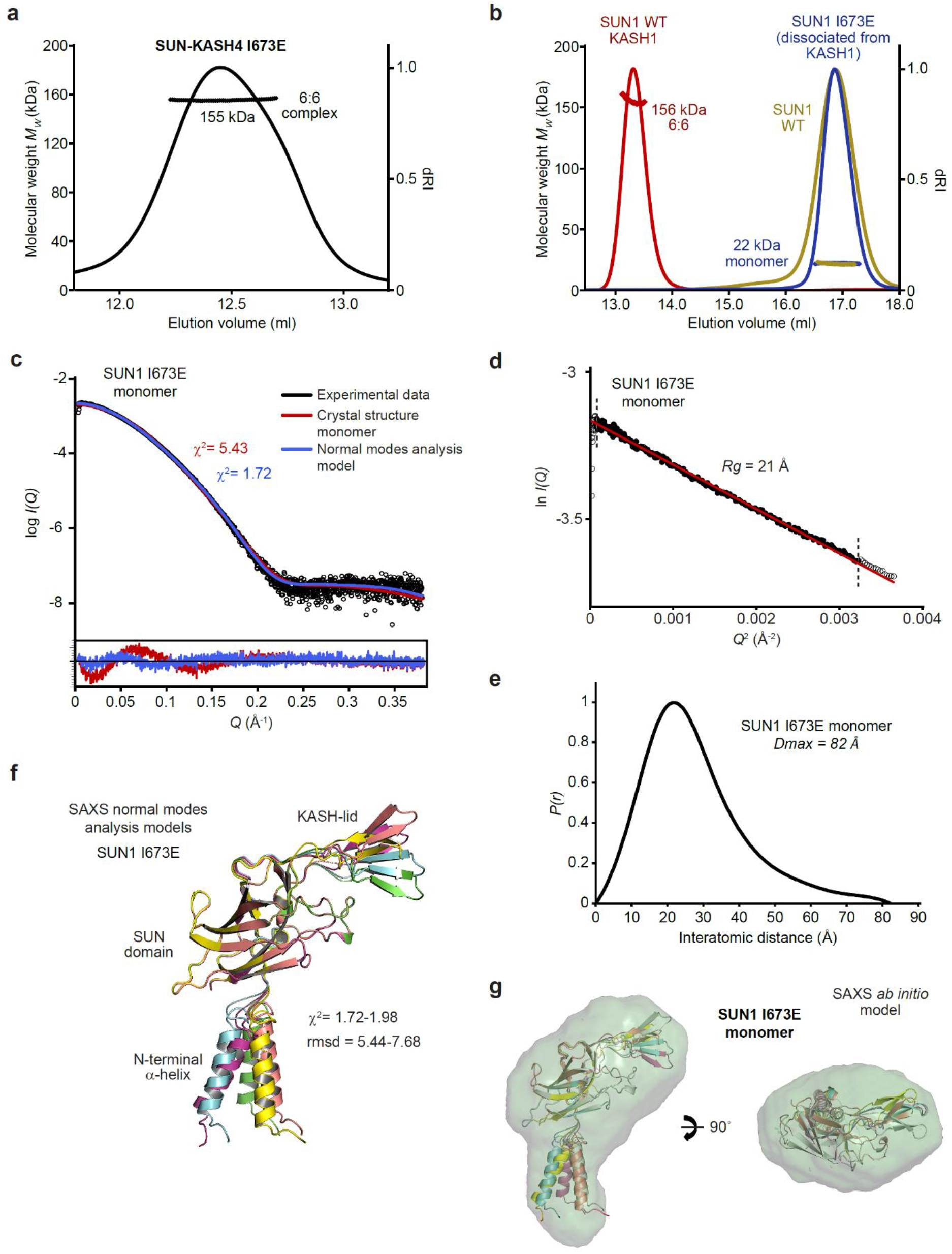
Biophysical analysis of the SUN1 I673E mutant. (**a**) SEC-MALS analysis demonstrating that SUN1 I673E and KASH4 form a 155 kDa 6:6 complex (theoretical 6:6 - 155 kDa). (**b**) SEC-MALS analysis demonstrating that isolated SUN1 wild-type (WT; yellow), and SUN1 I673E that has dissociated following co-expression with KASH1 (blue), are 22 kDa monomers (theoretical – 23 kDa). The wild-type SUN1-KASH1 6:6 complex is shown in red for comparison. (**c-g**) SAXS analysis of the SUN1 I673E monomer. (**c**) SAXS scattering curve overlaid with theoretical scattering curves of a SUN1 protomer from the SUN1-KASH1 crystal structure (red; χ^2^ = 5.43) and its normal mode analysis model (blue; χ^2^ = 1.72). (**d**) SAXS Guinier analysis revealing a radius of gyration (*Rg*) of 21 Å; the linear fit is shown in black and demarcated by dashed vertical lines (Q.Rg values were < 1.3). (**e**) SAXS *P*(*r*) distribution showing a maximum dimension of 82 Å. (**f**) SAXS normal mode analysis in which the five highest scoring models are displayed, based on their fit to the experimental SAXS data (χ^2^ = 1.72-1.98), and demonstrate hinge-like motion of the N-terminal α-helix relative to the SUN domain. (**g**) SAXS *ab initio* model (NSD = 0.645; reference model χ^2^ = 1.85) into which the normal mode analysis models are docked.

**Supplementary Figure 6.**
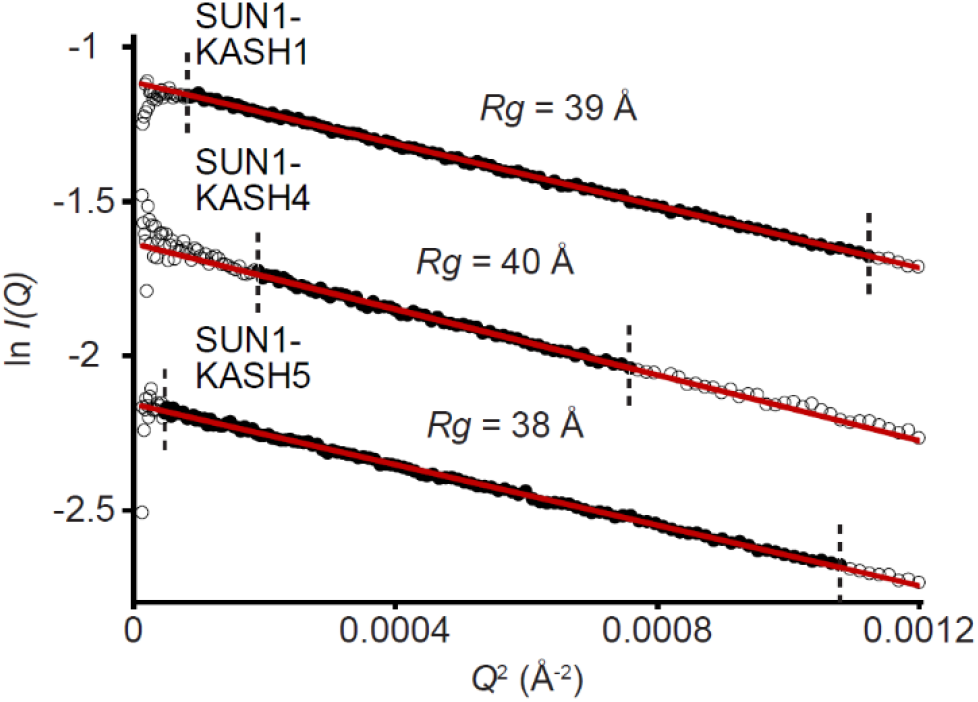
SAXS analysis of SUN-KASH complexes. SAXS Guinier analysis revealing radius of gyration (*Rg*) values of 38 Å, 40 Å and 39 Å, respectively; the linear fits are shown in black and demarcated by dashed vertical lines (*Q.Rg* values were < 1.3).

